# Neural encoding of pain uncertainty is selectively amplified when inferring another’s pain

**DOI:** 10.1101/2025.07.21.666065

**Authors:** L. Loued-Khenissi, G. A. Bergmann, C. Corradi-Dell’Acqua

**Affiliations:** Department of Clinical Neurosciences, Lausanne University Hospital, Lausanne University; Theory of Pain Laboratory, Department of Psychology, Faculty of Psychology and Educational Sciences (FPSE), University of Geneva, Geneva 1202, Switzerland; Department of Economics, University of Zurich; Center for Mind/Brain Sciences (CIMeC), University of Trento, 38068 Rovereto, Italy

## Abstract

Inferring another person’s pain is a challenging computational problem: unlike self-related pain, individuals have no access to others’ nociceptive input and must instead rely only on indirect cues under substantial uncertainty. Using a predictive-processing framework, we modeled how expectation, uncertainty (variance-based risk), prediction error (PE) and surprise jointly shape pain-related decisions for oneself and for a stranger. During fMRI, participants received cues signaling the intensity and probability of possible painful events, then made decisions to mitigate the anticipated pain via a lottery and a wager. Behaviorally, participants were more risk-averse toward others’ pain than their own, prioritizing pain reduction over economic gain. Neurally, anticipatory risk signals in anterior insula (AI) and dorsal striatum were selectively amplified when pain concerned another person, as opposed to self-pain. By contrast, PE and surprise were reliably represented in AIns for both targets following pain delivery. However, representational similarity analysis suggest that error-signal in AIns is represented through reliable, but dissociated, patterns for self and others. Together, these findings show that expectation, uncertainty and error processing are differentially affected by the identity of the pain recipient, and underscore how models of prediction under uncertainty are powerful tools for explaining how we assess pain in other people.

**Significance Statement:** When we make decisions about another person’s pain, we cannot directly access what they feel: we must infer it under uncertainty. Through a rigorous computational framework of prediction under uncertainty, we assessed the joint role played by expectancy, uncertainty and error-signal to model people’s decisions about their own and others’ pain. Deciding on another’s behalf increased brain and behavioral sensitivity to uncertainty; likewise error signals following a painful event were represented in a specific code, distinct from self-related pain. These findings offer a computational account of how humans navigate the inherent uncertainty of inferring others’ internal states, with direct relevance to clinical settings where surrogate decision-makers must estimate a patient’s pain despite lacking direct access to it.

---

Understanding pain is a challenging and uncertain inferential process, both when we experience it ourselves and even more so when it affects others. Self-related pain is not a direct readout of nociceptive input, but is shaped by expectation, its uncertainty, and deviation from prediction (1–9). In sharp contrast, a much scarcer literature has focused on the inferential processes underlying the assessment of another person’s pain, which cannot be accessed directly and must instead be extrapolated from indirect cues. The present study aims at modeling the inference of others’ pain within a unified computational framework of prediction under uncertainty, identifying associated neural correlates, and systematically comparing the extent to which results differ from those observed for self-directed pain.

Bayesian and predictive-processing accounts of the brain, posit that individuals build predictions about upcoming and compare them against actual outcomes (10–12). Predictions about outcomes are accompanied by representations of uncertainty (12) that refine subsequent estimates. The mean-variance approach formalizes this by expressing expectations as the average of the anticipated outcome distribution, and uncertainty as its spread, or variance (13–15). Uncertainty thus formalized is “risk”, the degree of unknown that can be anticipated in advance given known probabilities (16) (17). Comparing what was expected against what occurs yields two error signals: prediction error (PE), the discrepancy from the mean, and “Surprise”, the deviation from the variance (18). This approach offers a principled way of tracking the full inferential cycle: from anticipation, through the experience of uncertainty, to the updating that follows an outcome (14, 19, 20),

Absent nociceptive input, inference about others’ pain likely has consequences at every stage of the predictive cycle: expectations about others’ pain may be systematically biased, uncertainty amplified, and prediction errors may be weighted differently. These distortions are not merely theoretical: research has repeatedly shown that perceptions of others’ pain are shaped by contextual information (21–24) and social influence (25). In the clinic, healthcare practitioners show systematic biases in pain estimation and management, reflecting experience-dependent underestimation susceptibility to contextual information (26), and heightened sensitivity to uncertainty and error signals (27). Such misestimations could plausibly arise at any of several distinct stages: in the expectation itself, in its associated uncertainty, or in how new information updates estimates going forward.

For self-pain, expectancy effects are robust: cues predicting higher pain lead to stronger experiences, while those predicting relief produce hypoalgesia (placebo effects) (1). Uncertainty effects are less consistent: while some studies report enhanced pain following cues associated with high unpredictability or risk (1, 28, 29), but a meta-analysis did not confirm this effect (30) (see also (31). In lottery tasks, individuals generally prefer risky options when facing the prospect of negative events, such as economic loss, disease or pain (32–34), yet seem less willing to gamble for another’s pain than for their own; cue unpredictability likewise makes individuals less sensitive to others’ pain (34).

Neuroimaging offers a complementary route to this question, since numerous studies have documented a key role for the insular cortex in different aspects of self-pain expectancy and experience. The posterior insula processes nociceptive input (35, 36), whereas the anterior insula (AIns) plays a more active role in inference: middle-dorsal portions integrate nociceptive information with prior expectations while ventral-anterior portions underlie PE signals (29). Much less is known about the neural coding of uncertainty, though some work points to AIns (29) as well as parietal cortex (37). Abundant evidence implicates AIns in processing information about others undergoing this experience (38, 39). This dual sensitivity is usually read as a unitary process coding what the two targets share: empathy (40), unpleasantness (41), or salience (42–44). Yet, AIns has also been implicated in coding risk (14, 45), PE (46, 47) and surprise (18, 48), across several domains. A second structure of high theoretical relevance is the striatum, where distinct subregions have been linked to the processing of expectancy (14, 49), risk (14) and PE (46, 47). A striatal role in expectancy and predictive coding has also been documented in pain research, chiefly in meta-analyses of placebo effects (50). Both these regions are strong candidates for supporting each stage of pain-related inference. However, whether they track expectation, uncertainty, and prediction error as dissociable signals within a single pain paradigm, and whether this tracking differs for self *vs.* other, has yet not been tested.

In this study, we examined how a computational framework of prediction under uncertainty can explain pain processing for both self and others. Participants completed an fMRI task where predictive cues signaled the intensity and probability of possible painful events affecting either themselves or a stranger, and made decisions to mitigate the anticipated pain via a lottery and, subsequently, a wager (Figure 1A-B). Modeling yielded trial-wise estimates of average Expected Pain (EP; intensity weighted by probability), Risk (the variance across possible outcomes), and, following pain delivery, signed PE and Surprise (Figure 1C).

**Figure 1.**
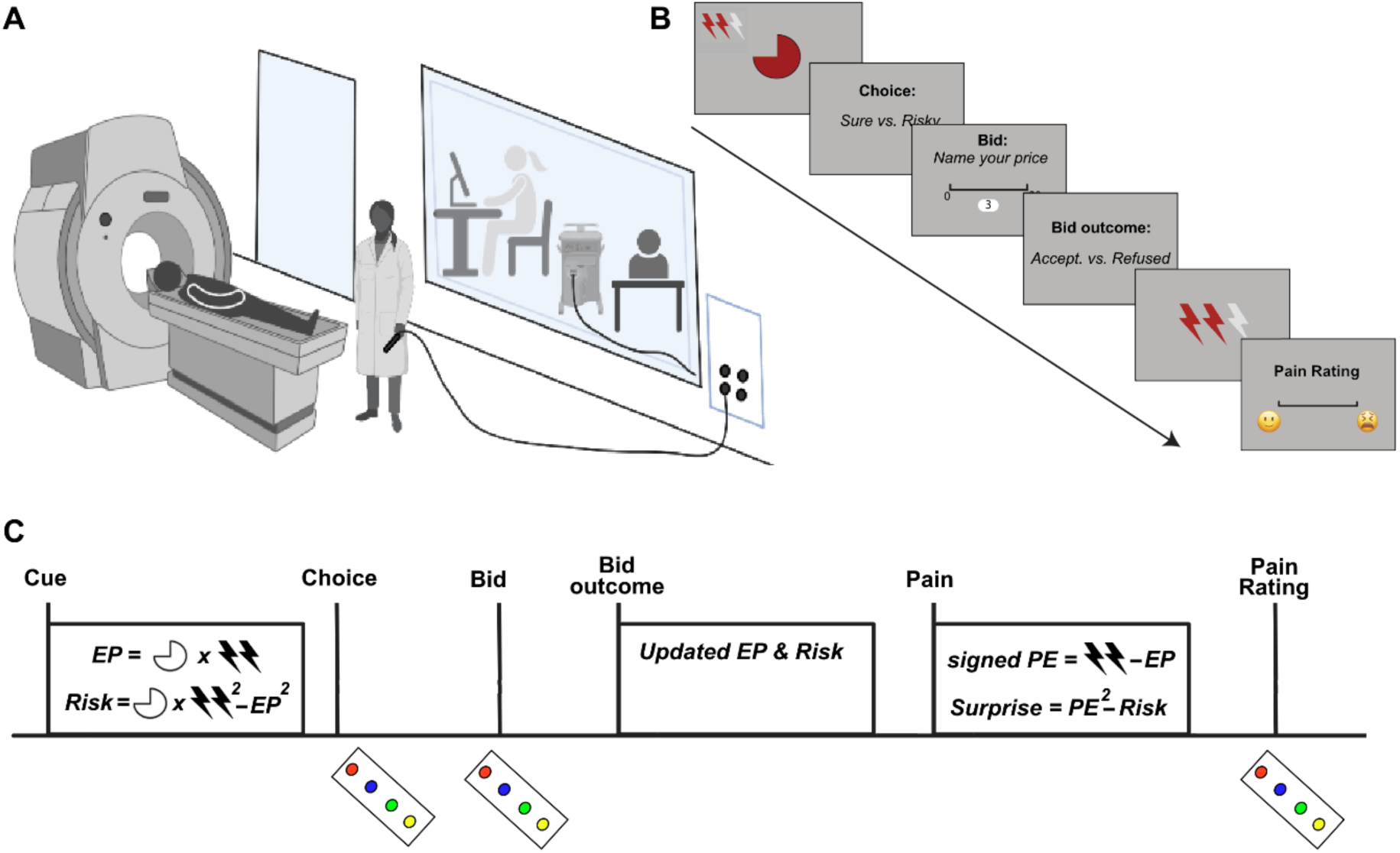
A) Experimental setup. Deciding agents made decisions on threatened pain for themselves and others during an fMRI session. An experimenter delivered pain via laser within the scanner room. The participant was made to believe the other remained in the control room and was subject to pain delivery during the “Other” block condition. B) Sample Trial. Participants were first shown a visual cue representing an initial value of expected pain with the probability of delivery exhibited in a pie chart, and pain intensity by red lightning bolts. They were then asked to 1) either gamble to avoid pain or; 2) choose a sure option to reduce pain. They were then asked to place a monetary bid on their choice, which was then compared to a hidden, randomly generated reserve price. The participant was then told whether the bid was accepted or not, and the pain outcome was subsequently shown or delivered based on the expected value of the threatened pain following bid outcome. Participants were then asked to rate the perceived pain, for themselves in the “Self” condition, or the other felt in the related condition. C) fMRI Design. The general linear model constructed for fMRI analysis included the following epochs: cue onsets, choice onsets, bid, bid outcome, pain event and pain rating. Trial-wise estimates of Expected Pain (EP) and Risk were estimated from the anticipatory phase prior pain delivery, whereas signed Prediction Error (PE) and surprise were estimated following the painful event.

We expected strong modulation of AIns and the striatum at each predictive stage for self-pain. The critical question is whether the same holds for the assessment of others’ pain. This will allow us to determine the extent to which expectation, uncertainty, PE, and Surprise processing are equally, or differentially, affected by the identity of the social recipient.

## Results

Of 36 participants, 3 were excluded (2 for calibration errors, 1 for coil misplacement), leaving 33 deciding agents (13 male; mean age 23.2 ± 4.3 years) in 33 dyads.

### Behavioral Results

As preliminary step, we assessed whether, and to which extent, participants’ representation of probabilities was distorted relative to their objective values. A Prelec probability weighting model (51) based on individual wagers offered for pain relief (hereafter, bids) yielded subjective values that were then compared to objective task-derived regressors, confirming that the degree of distortion was trivial, and the manipulated probabilities were indeed appropriate for the analysis (Figure 2A; see also Supplementary Information S1 for more details).

**Figure 2.**
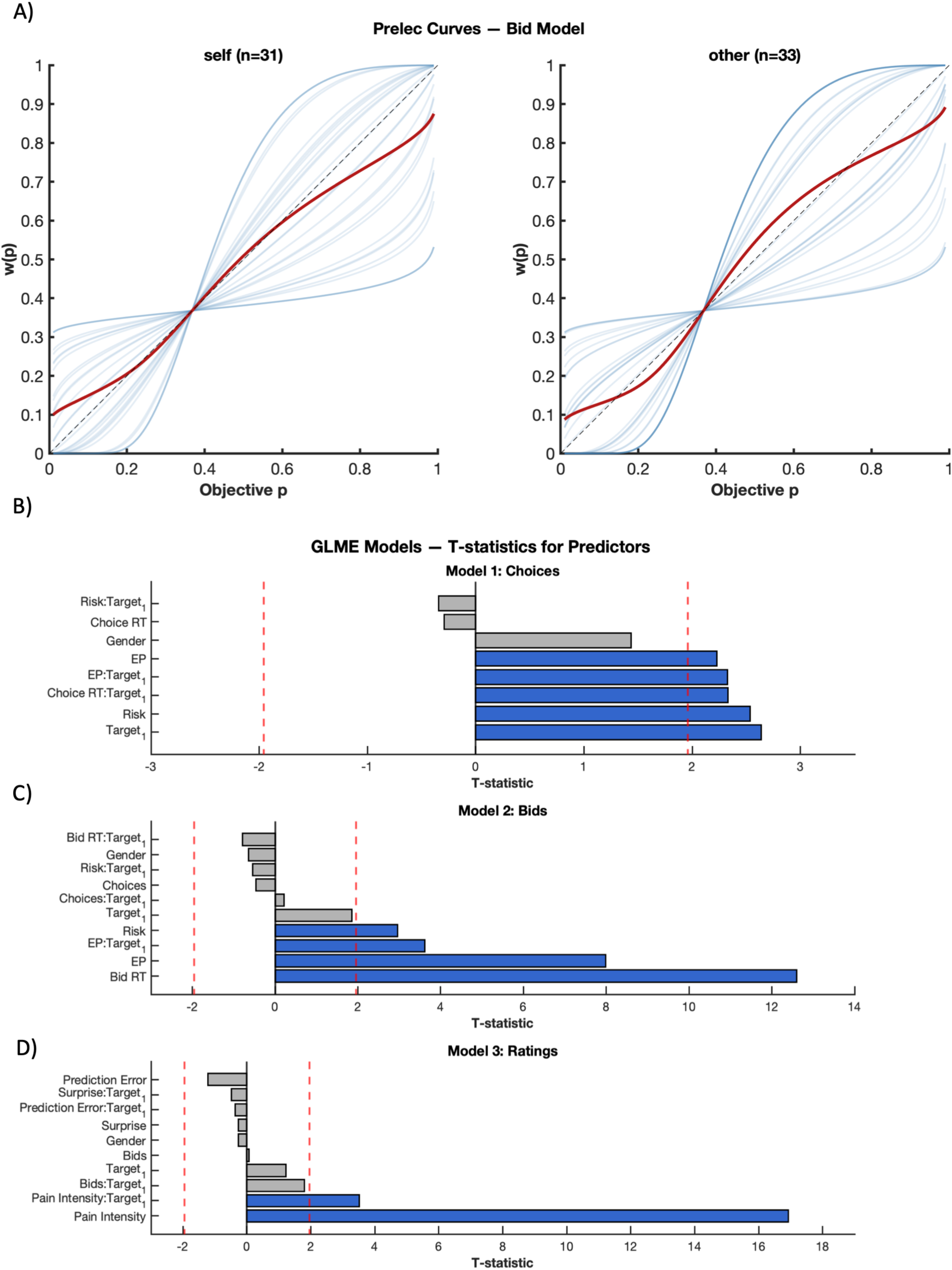
Behavioral results and computational modeling of pain inference. A) Subjective probability weighting (Prelec). Estimated Prelec functions mapping objective probabilities to subjective decision weights for Self (n=31) and Other conditions (n = 33). Two participants in the Self condition placed 0 bids at every trial and could not be included in the Prelec function. T-statistics for fixed-effect predictors from three (generalized) linear mixed-effects (GLME) models: (B) choice behavior, (C) bid value, and (D) pain intensity ratings. Each horizontal bar reflects the t-statistic for a fixed-effect term; red dashed lines mark approximate significance thresholds (|t| > 1.96). The x-axis is scaled dynamically to reflect the range of t-values, with the left bound fixed at –3.

Subsequently, to assess how computational variables influenced behavior, we constructed three mixed-effects models targeting distinct stages of the decision process. Model 1 tested whether EP, Risk, and target identity predicted choice (gamble vs. sure option) modeled as a dichotomic variable. Model 2 tested whether EP, risk, choice, and target predicted bids for pain relief. Model 3 tested whether pain intensity, signed prediction error, surprise, and target predicted subjective pain ratings. Overall, all three models revealed a consistent pattern of heightened risk aversion for others. Expected pain (t = 2.23, p = .026), risk (t = 2.54, p = .011), and target identity (t = 2.64, p = .008) predicted a higher likelihood of sure-option selection in Model 1. Likewise, in Model 2, higher bids were associated with higher EP (t = 7.98, p < .001) and risk (t = 2.96, p = .003); furthermore, EP effects were stronger for others, as revealed by a EP × target interaction (t = 3.62, p = .003). Finally, Model 3 revealed that pain ratings were on overall higher for others’ pain (t = 3.51, p = .005), with no effect of signed prediction error or surprise on ratings (Figure 2B-D).

Taken together, results suggest a differential pattern of choice behavior for a pain threat to another, in the form of heightened risk aversion in both Models 1 and 2, higher pain relief valuation for the other in Model 2, and inflated pain ratings in Model 3 (Figure 2D). A direct effect of uncertainty (risk) on decision-making was only found in the prediction phase; neither prediction errors affected subjective pain ratings.

### Neuroimaging Results

#### Expected Pain and Risk modulate others’ related pain anticipatory activity

Full univariate neuroimaging results are listed in Supplementary Information S3. While looking at neural responses associated with the task, we distinguished between the pain anticipatory phase (corresponding to the epochs where choices/bids were made) from the painful outcomes. During the anticipatory phases, trial-wise estimates of EP and Risk were modeled as parametric modulators to assess for brain regions that kept track of these parameters’ fluctuations. For both EP and Risk, suprathreshold effects were observed only when pain targets others (cluster-corrected FWE threshold corresponding to p < 0.05, cluster-forming height threshold corresponding to *p* < 0.001 uncorrected; see methods). EP effects of others were associated with enhanced activity in the visual cortex, extending to the fusiform gyrus; as well as to the supplementary motor area (Figure 3, in blue). Risk effect for others was associated with widespread activity that spanned occipital, temporal, middle cingulate, striatal, insular regions as well as cerebellar responses (Figure 3, in green). No suprathreshold effect was observed for self-related anticipatory phase. Furthermore, direct comparison between EP effects in Others > Self confirmed the role of the left Fusiform gyrus. Similar comparison of Risk effects in Others > Self implicated the middle temporal gyrus.

**Figure 3.**
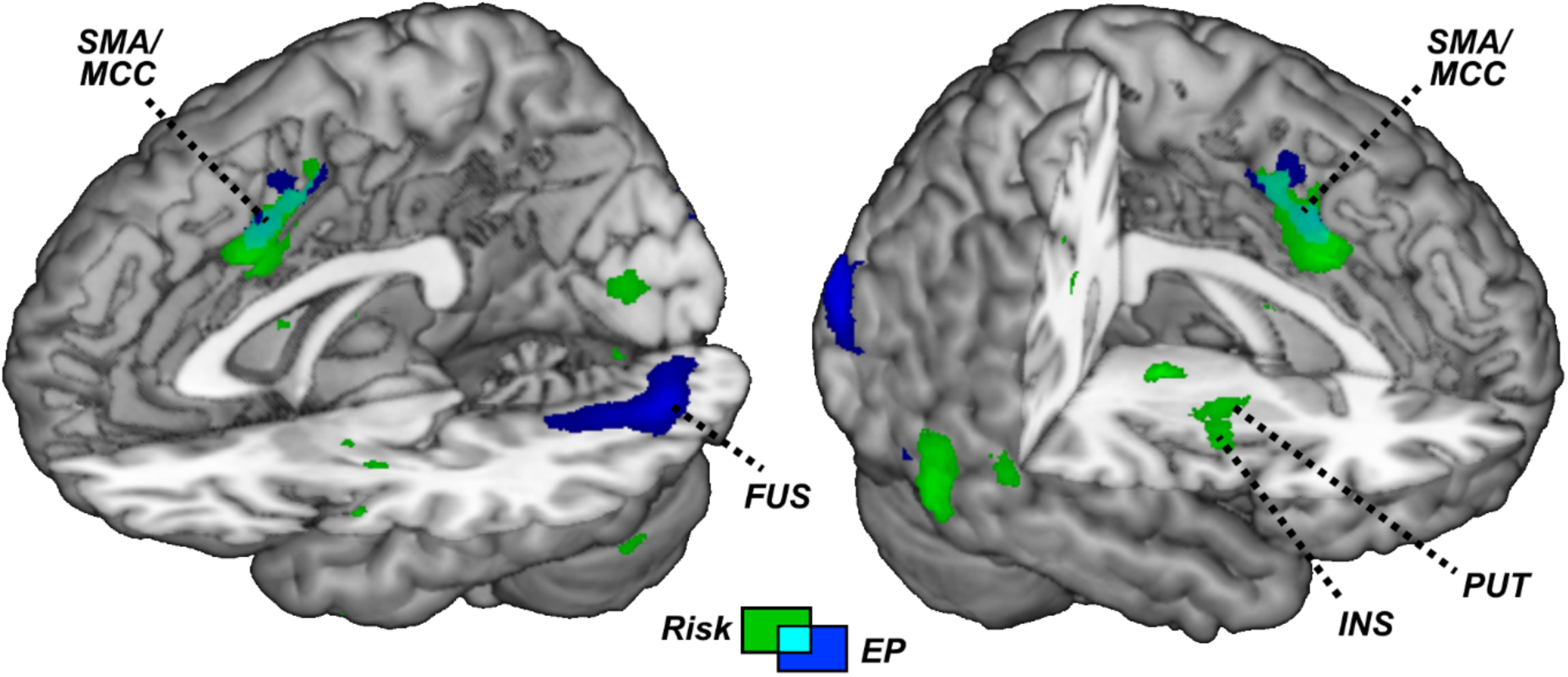
Brain surface rending and activation maps displaying neural structures responding to Expected Pain (blue blobs) and Risk (green blobs) during the anticipation of others’ pain. SMA = Supplementary Motor Area; MCC = Middle Cingulate Cortex; FUS = Fusiform Gyrus; INS = Insula; PUT = Putamen.

We complemented whole brain analysis with targeted Region of Interest (ROIs) analysis for six anatomical regions of interest: anterior insula, caudate and putamen in each hemisphere. Small-volume FWE correction analysis revealed significant effects only for Risk. All six ROIs were associated with a reliable anticipation of others’ pain Risk, whereas only the left caudate was associated with anticipation of one’s own pain Risk.

Furthermore, right AIns and Putamen showed risk effects reliably higher for Others than for self (see supplementary Table S1.1 for full ROI analysis).

We also performed correlation-based Representation Similarity Analysis (RSA) to establish distributed patterns of activity within each ROI that represented information about either EP or Risk. Under correction for multiple comparison for the six ROIs no significant effect was found.

#### Self and other activity associated with the pain event

We then looked at the activity associated with the painful event. Figure 4A provides information about the neural structures implicated in pain intensity. Consistently with a wide literature documenting shared activity between one’s own and others’ pain (38, 39), we confirmed a key role of right anterior insula (Figure 4, in yellow) in processing pain intensity for both self (as identified through whole brain analysis) and others (through small volume correction). Self-related pain experience was also observed in a widespread network encompassing the whole insular cortex, middle/anterior cingulate cortex, dorsolateral prefrontal cortex, etc. (in red), while other-related pain triggered activity in the occipital cortex (in green).

**Figure 4.**
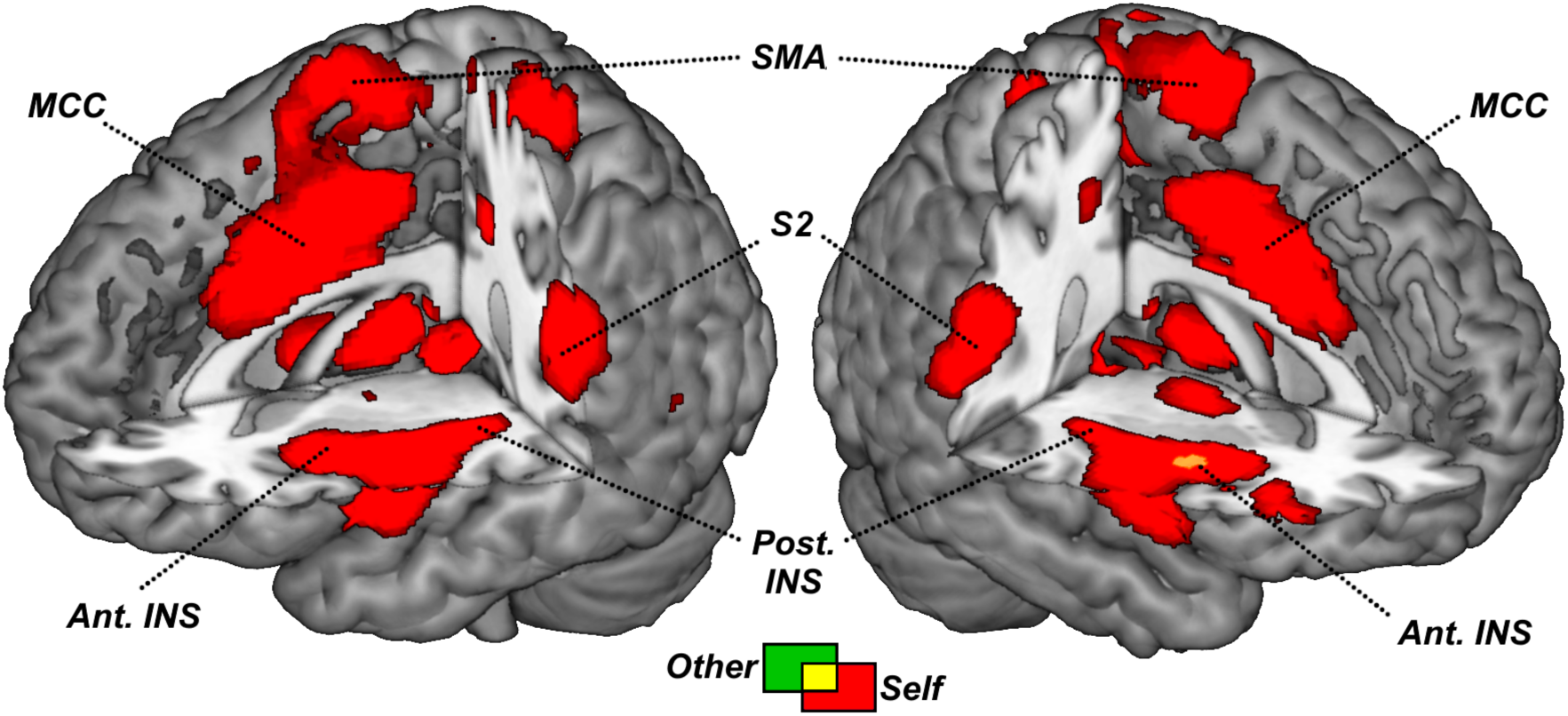
Brain surface rending and activation maps displaying neural structures parametrically associated with the intensity of Pain outcome when delivered to participants’ themselves (red blobs) or an unknown stranger (green blobs). SMA = Supplementary Motor Area; MCC = Middle Cingulate Cortex; Ant. & Pos. INS = Anterior & Posterior Insula; S2 = Secondary Somatosensory Cortex

We looked at the activity evoked by signed PE and surprise. No effect was associated with signed PE, neither under rigorous whole brain correction for multiple comparison, nor under ROI analysis. However, surprise for one’s own pain revealed activity in the Postcentral Gyrus (Figure 5). ROI analysis for surprise was carried out within the *a priori* domain-general surprise region of anterior insula, defined independently from perceptual and economic surprise (18). Small-volume correction revealed significant surprise responses within this region for both self (T = 3.90, p(FWE) = .016) and other (T = 3.43, p(FWE) = .039), with no significant self-vs-other difference.

**Figure 5.**
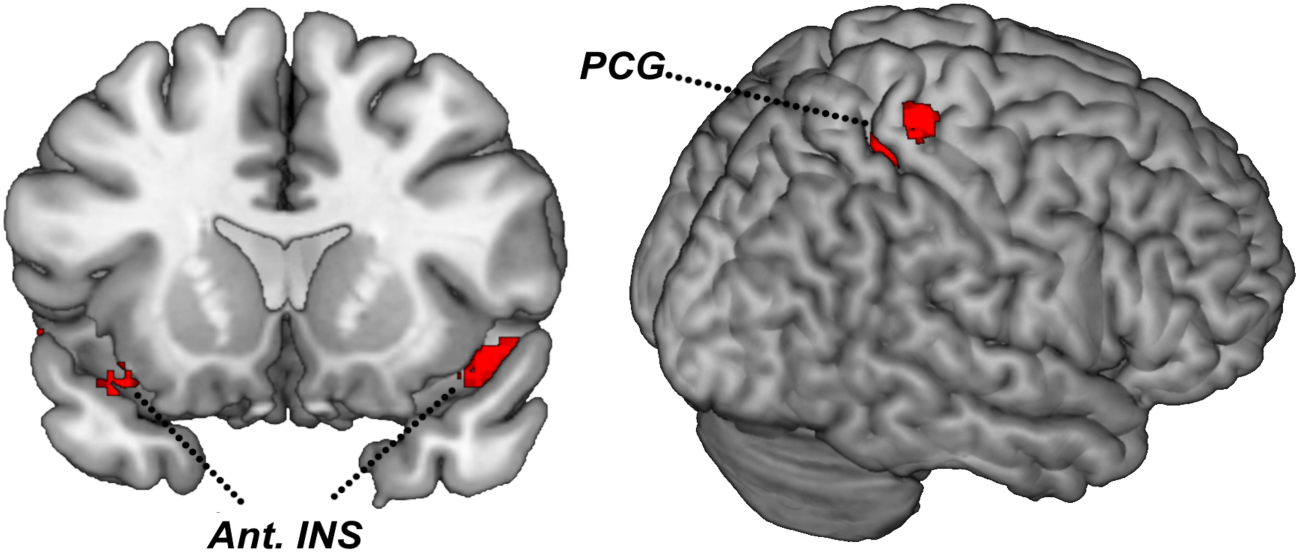
Brain surface rending and activation maps displaying neural structures parametrically associated with pain Surprise in the self-condition. PCG = Post-Central Gyrus; Ant. INS = Anterior Insula.

In addition to standard univariate analysis, we performed RSA to identify distributed regional patterns that represent PE and surprise. Bilateral anterior insula carried significant representational information on signed PE for *both* self and other (see Table 1 & Figure 6). The representational geometry of pain prediction error was, however, different across targets. By pooling together trials from the self- and other-conditions, we assessed the degree of representational similarity between trials of the same target (within-target similarity) relative to trials of different targets (cross-target similarity). For both right and left AIns significant representational information was observed for within-target similarity, but not cross-target similarity. Furthermore, within-target similarity parameters were systematically higher than those associated with cross-target parameters (Figure 6). This reveals that AIns represents prediction error for both self and others’ pain, but this information appears to be target-specific. Interestingly, a similar pattern was also observed in the right Caudate, but the effect did not survive correction across the 6 ROIs for the self-condition (see Table 1).

**Figure 6.**
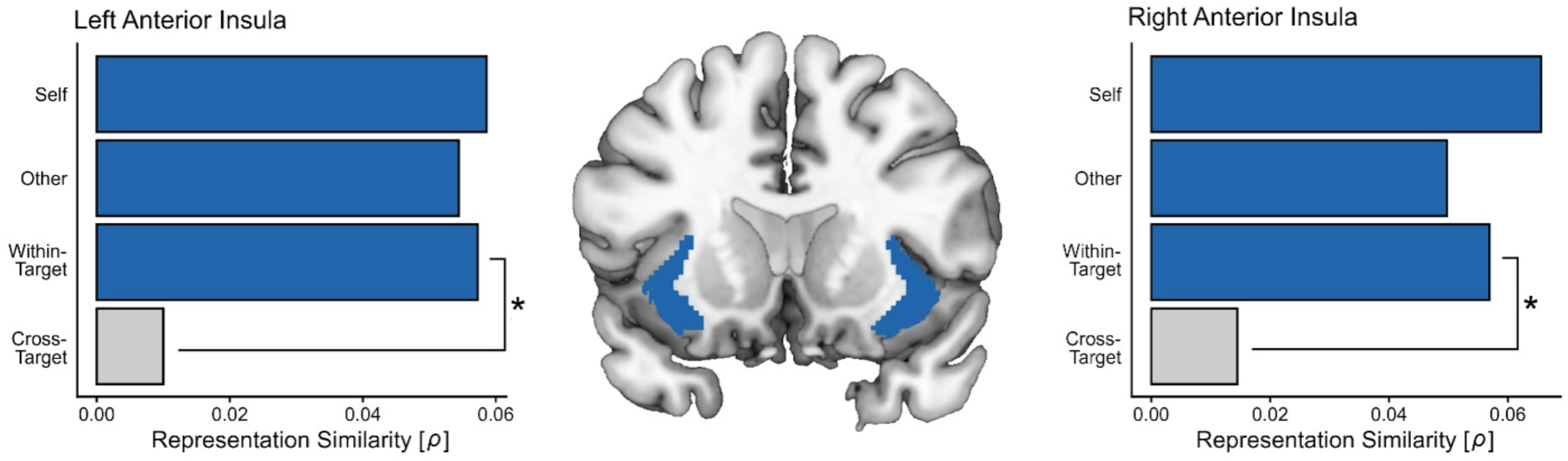
Representational similarity analysis for signed prediction error in the anterior insula (ROI) analysis. Bars represent Spearman correlations between neural and model dissimilarity. Blue bars represent effects significantly higher than 0 under FDR correction for multiple comparisons for the 6 ROIs. “*” represent significant differences between within-vs. cross-target representation similarity under FDR correction.

**Table 1.** Representational similarity analysis (RSA) for signed prediction error. Values represent Spearman correlations between neural and model dissimilarity. For within-vs. cross-target comparisons values represent differential spearman ρ correlation coefficients.

| Region | Self | Other | Within-T | Cross-T | Within vs. Cross |
| --- | --- | --- | --- | --- | --- |
| L Anterior Insula | .059* | .054* | .057* | .010 | .047* |
| R Anterior Insula | .066* | .050* | .057* | .014 | .042* |
| R Caudate | .040^ | .048* | .048* | .012 | .036* |
| * p < 0.05 FDR corrected for the 6 ROIs; ^ p < 0.05 uncorrected |  |  |  |  |  |

## Discussion

By exploiting a computational framework of prediction under uncertainty, we dissected the components underlying inferences about other people’s pain, and identified similarities and discrepancies with the assessment of one’s own pain experience. At the behavioral level, participants were more risk-averse toward others’ pain than toward their own, as shown by both choice and bidding behavior. This was mirrored by positive BOLD responses during the anticipatory epoch preceding pain delivery, for risk of others’ pain in a widespread network including AIns and striatum, an effect not observed for self-pain. By contrast, analysis of the painful events themselves revealed reliable effects of PE and Surprise for both targets. In particular, Representational Similarity Analysis (RSA) confirmed that AIns represents signed PE information for both one’s own and others’ pain, although the representational geometry differed between targets, pointing to target-specific coding. Together, these findings shed new light on processes underlying pain expectancy and uncertainty, and underscore how models of prediction under uncertainty are powerful tools for explaining how we assess pain in other people.

### Risky decision-making for others

Our behavioral results largely replicate previous findings, revealing a consistent pattern of risk aversion when deciding for others (34), in contrast to reports of equal or increased risk-taking in other-regarding monetary decisions (52). As in our earlier work, we observed a higher monetary valuation of others’ pain relief compared to one’s own (34), suggesting a cautious approach to vicarious pain evaluation. More risk-averse, costlier decisions may reflect altruistic motivations reported in other studies (53–55) but could equally reflect a drive to minimize the uncertainty introduced by the decision-maker’s lack of direct access to the target’s nociceptive state. Taken together, the discrepancies between self- and other-related behavioral effects may reflect differences in uncertainty weighting within the same predictive-processing framework, rather than fundamentally distinct decision processes.

### Neural representation of risk and expectancy in others

In the other condition, we found significant neural responses to pain risk that were decoupled from pain expectation, in both AIns and the dorsal striatum, a region reliably implicated in decision-making under risk (56). Striatal and insular responses to pain risk arose when decisions concerned others, consistent with the behavioral risk aversion observed in vicarious contexts. This selective response is compatible with the interpretation that, lacking direct sensory access to the other’s nociceptive state, the brain overweights uncertainty computations when inferring another’s pain, perhaps by assigning a decreased precision to a prediction (57, 58). Expected pain for others additionally recruited visual regions, including the fusiform gyrus, which may reflect enhanced attentional allocation toward cues predicting another’s pain outcome (59, 60). The fusiform gyrus is frequently implicated in processing facial information, with heightened activity when faces convey emotional information (see (61)as review). It is therefore conceivable that cues predicting high EP triggered an anticipatory representation of the other person’s reaction, including their expected facial response. Future work will need to address this possibility directly.

### Neural representation of error signals in self and other

In contrast to the target asymmetry observed for risk, error-related representations were reliably found in both targets. Although univariate analysis of signed PE yielded no significant effects, complementary RSA revealed a significant pattern of AIns activity tracking trial-by-trial PE changes in both self and other conditions. Critically, the fact that AIns represents PE for both targets does not necessarily indicate a single, shared process for error estimation: direct comparison of within-target *vs*. cross-target similarity showed that self- and other-related error signals are characterized by distinct representational geometries within the insular cortex. Previous research has repeatedly documented shared activity patterns within AIns for self- and other-pain (26, 62–66), generalizing even to other aversive experiences such as disgust and unfairness (41, 63). Our findings suggest that this shared activation does not imply a shared error representation, as PE effects in our data are clearly target-specific. Our data already reveled target differences in the processing of anticipatory EP and Risk, suggesting that target-selective weighting of anticipatory information may extend to a similarly target-selective comparison of outcomes against expectations.

Prior work comparing expectancy for self-directed pain against other aversive events (e.g., disgust, shocking images) found that AIns can run parallel, domain-specific predictive models for different classes of negative events (35, 67, 68). Our results converge with, but also extend, this work, pointing to an inferential architecture capable of running concurrent but separate predictions depending on the pain recipient.

### Surprise implicates AIns in both self and others

However, in contrast to signed PE, Surprise effects were captured only by univariate analysis and not by RSA, making it difficult to determine conclusively whether self- and other-related effects share a common representational code or not. In principle, RSA (which relies on the joint activity of multiple coordinates at once) should capture information beyond the reach of univariate approaches. However, RSA is sensitive to pattern variability rather than overall signal magnitude, and may therefore miss critical information that drives univariate effects. Keeping these considerations in mind, our surprise findings converge with prior work showing that similar coordinates respond to surprise across perceptual and economic judgments (18), suggesting the presence of a computational AIns mechanism that signals departures from expected uncertainty across task domains. The definition of surprise used here (i.e., events that exceed the expected spread of outcomes, see Introduction) is also reminiscent of the concept of perceptual salience in neuroimaging research, where part of the insular signal (along with other regions) is thought to respond to events that stand out relative to their context, capture exogenous attention, and promote behavioral adaptation (44). While prior studies have typically manipulated salience through abrupt sensory changes (e.g., a bright light, a loud noise, nociceptive input) (44), it is plausible that similar circuitry identifies unforeseen departures from prior expectations, prompting adaptation of ongoing predictive models. Just as insular salience coding is thought to operate jointly with pain-specific information (44), surprise coding in our study may similarly integrate target-specific information (risk and PE), contributing to overall pain assessment.

### Limitations

Our study carries limitations that should be acknowledged. First, consistent with our prior work using similar paradigms (33, 34), we found evidence of costly other-regarding tendencies, with participants consistently prioritizing others’ pain over economic gain. This may reflect an altruistic desire to reduce the other’s pain (69) (70, 71), but could equally stem from reputational concerns tied to being observed (the so-called “experimenter effect” Hoffman et al., 1996). Second, our paradigm did not provide agents with cues such as the target’s facial expressions (73) or vocalizations (74) unlike our earlier work, where agent and target were seated together (75). This difference in context may account for the behavioral discrepancies between the present and prior study.

## Conclusions

This study provides evidence for a neurally instantiated, graded representation of another’s threatened pain. We show that both shared and target-specific computations of expectation, uncertainty, prediction error, and surprise contribute to how we assess the pain of another person. This work extends the applicability of computational models of prediction under uncertainty to how we navigate another’s suffering, with implications for clinical contexts where surrogate decision-makers must act under irreducible uncertainty about another’s pain.

## Materials and Methods

### Population

A total of 72 participants (36 dyads) were enrolled in the study. Sample size was comparable to prior model-based fMRI studies on uncertainty (n≈25–35). Participants were recruited from both the University of Geneva Campus and the general population. Inclusion criteria included: French-speaking; 18 -45 years of age. Exclusion criteria included: reported psychiatric or neurological diagnoses; current or recent psychotropic drug use; MRI incompatibility. Participants were randomly assigned a fixed role in the task: they were either cast as a Deciding Agent (DA), or a Passive Recipient (PR). The DA’s role in the experiment was to enter the scanner, and make decisions for both themselves and the PR. The PR’s role was to act as the recipient of half of the DA’s decisions. Three DAs were excluded from analysis (two for misapplied laser fluence, one for coil misplacement), leaving 33 (13 male; mean age 23.22, SD 4.33). The study was approved by the Commission Cantonal d’Ethique de la Recherche sur L’Etre Humain in December, 2019.

### Procedure

DAs were led to believe that PRs were undergoing a pain calibration procedure and would receive stimulation on half of all trials. In reality, no such procedure was performed, as laser pain stimulation could not be administered in the control room due to safety constraints related to glass walls at the center. A deception was introduced whereby DAs believed that PRs received pain on half of the task trials. After DAs were positioned in the scanner, PRs were paid 30 CHF, debriefed and dismissed. DAs were informed at the end of the experiment that PRs had not received pain; all DAs believed that a real person was present and potentially affected by their decisions.

### Pain Calibration Procedure

Nociceptive radiant heat was delivered to the ankle via an Nd:YAP laser (Stimul 1340, El.En, Florence; 1340 nm, 4 ms pulse, 6 mm² beam). Energies from 1.5–3 J were emitted in random order at 0.25 J steps, three times each, and rated on a 0–10 scale (75). Averaged ratings closest to 1, 5, and 9 defined subjective low, medium, and high levels used in the task.

### Task

DAs performed two blocks of 18 trials of a decision-making task, one targeting themselves and one targeting the partner. At each block, participants started off with an endowment of 216 monetary units (MUs), equivalent to 21.6 CHF. At each trial, DAs were first presented with a visual cue, depicting a probability (in the form of a pie chart), and an intensity (represented by 2 or 3 red lightning bolts) of threatened pain, for 6 seconds (Figure 1B). Probabilities ranged from .1-0.9 in steps of .1. DAs could then either gamble to avoid the pain; or select a sure option, where pain would be reduced by one level. After choice, they bid on their selection against a hidden reserve price. Accepted bids applied the choice; rejected ones left the original threat standing. Pain was delivered depending on resultant probabilities and intensities. A visual cue representing this pain delivery was simultaneously presented on the in-bore screen, as either initial values of pain intensity (2 or 3 red lightning bolts); reduced values of pain intensity (2 or 3 yellow lightning bolts) or no pain (3 greyed out lightning bolts). Finally, DAs were asked to rate the pain outcome on a visual rating scale (Figure 1B).

### Computational model

Trial-wise expected pain (EP = *p* × *P*) and risk (σ² = *p* × *P*² − EP²) were derived at cue onset and updated after each bid outcome. Following trial outcome, both signed prediction error (PE = observed − EP) and Surprise (PE² − σ²)(76), were computed. A one-parameter Prelec weighting function (51) fitted to bids yielded the subjective counterparts of these values ; equations and fitting are given in *SI Appendix*, S1 and S2.

### Behavioral analysis

Choices were modeled with a binomial mixed model, bids and ratings with general linear models, each including the relevant computational variables, reaction time, and their interactions with target, plus gender as a nuisance factor and participant as a random effect.

### fMRI acquisition and analysis

Whole-brain echo-planar volumes were acquired on a 3T Siemens scanner (TR 1100 ms, TE 32 ms, flip angle 50°, multiband) alongside a T1weighted structural scan. Preprocessing in SPM12 proceeded as follows: slice-time correction, distortion correction, realignment, coregistration, normalization to MNI152, and 8 mm FWHM smoothing. First-level models convolved onsets with the canonical hemodynamic response and included parametric modulators for each computational quantity above, as well as 24 physiological and motion nuisance regressors, and FAST autocorrelation control. Group-level one-sample *t* tests were thresholded at cluster-level *p* < 0.05 FWE (voxelwise *p* < 0.001). Region-based analyses focused on AAL3 (76) masks for anterior insula, caudate, and putamen; prediction error contrasts were tested via an insula mask defined functionally from a previous study (18).

### Representational similarity analysis

Neural dissimilarity matrices (1 − Pearson *r* across trial-wise patterns) were compared using Spearman ρ against model matrices for EP, Risk, PE, and Surprise, within and across targets. Significance was assessed by permutation (10,000) and FDR correction across regions.

## Acknowledgments

This work was supported by Swiss National Science Foundation grant PP00P1_183715 awarded to C.C.D. This study was supported by the MRI Platform, **neuro** @campusbiotech, Geneva, Switzerland, co-founded and supported by Ecole Polytechnique Fédérale de Lausanne (EPFL), University of Geneva (UNIGE), and Geneva University Hospitals (HUG).

## Data Availability

Neuroimaging data have been deposited in OSF (https://osf.io/da7tw) and behavioral data in OSF (https://osf.io/uqtcw). Analysis code is available at https://github.com/LLouedKhen/OPPI_Project_Unige2021. All other study data are included in the article and/or *SI Appendix*.

## Author Contributions

L.L.K. and C.C.D. designed research; L.L.K. and G.A.B performed research; L.L.K. and C.C.D analyzed data; and L.L.K., G.A.B., and C.C.D. wrote the paper.

## Competing Interest Statement

The authors declare no competing interest.

## Notes

### Competing Interest Statement

The authors have declared no competing interest.

### Summary of Updates

This version has refined analyses relating to the neuroimaging portion of the data, and added an additional analytical check using RSA.

