## Supplementary information for "Neural encoding of pain uncertainty is selectively amplified when inferring another’s pain"

#### S1. Methods

##### *Population*

The study was conducted at the Campus Biotech in Geneva, Switzerland. A total of 72 participants (36 dyads) enrolled in the study. Sample size was comparable to prior model-based fMRI studies on uncertainty (n≈25–35). Participants were recruited from both the University of Geneva Campus and the general population. Inclusion criteria included: French-speaking; 18 -45 years of age; and good general health. Exclusion criteria included: reported psychiatric or neurological diagnoses; current or recent psychotropic drug use; claustrophobia; metal or other implants that may interfere with the MR scanner. Study information, including the task procedure and MR safety guidelines, was sent electronically to all participants 2 days before the scheduled experiment session. Upon arrival at the study center, participants were given paper versions of the information and consent forms. Participants were asked to read these, confirm their understanding of the study's risks and benefits, and sign the forms before continuing with the experiment. Following intake, the experimenter verbally explained the study structure and task process. Once participants understood the experiment, they were each randomly assigned a fixed role in the task: they were either cast as a Deciding Agent (DA), or a Passive Recipient (PR). The DA's role in the experiment was to enter the scanner, and make decisions both for herself and the PR. The PR's role was to act as the recipient of half of the DA's decisions. The study was approved by the Commission Cantonal d'Ethique de la Recherche sur L'Etre Humain in December, 2019. Data collection was conducted between April and July, 2021.

Of the 36 participants enrolled as deciding agents (DAs) in the study, 2 were excluded from analysis because laser fluence levels for the hand (which are lower in energy range) were erroneously used on their ankles, resulting in a lack of perceived pain. An additional participant was excluded from analysis due to a misplacement of the MR head coil, resulting in only half of the coils receiving signal. The remaining participant pool comprised 33 DAs (13 Male; average age, 23.22 years, sd = 4.33) participating in 33 dyads. Efforts were made to counterbalance DA sex when possible, alternating between males and females, but the participant pool consisted mostly of women. Twelve dyads consisted of agents of different sex across the dyad. Mean payout was 17.28 CHF (sd = 3.65). Of the 33 participants, 5 consistently made the same choice for themselves across trials; of these, 3 always gambled and 2 always selected the sure option. Two participants (one of which was also one of the 5 consistent self gambles) always selected the same option for the other (1 gambler and 1 fully risk averse). Finally, 2 participants made no bids in the self condition. All participants bid > 0 CHF for the other condition.

##### *Procedure*

Following intake, both participants were asked to remove metallic items and leave personal affairs in dedicated lockers before being led to the scanner control room. Here, both participants were briefly shown the laser pain stimulation machine and the PR was asked to take a seat in the control room. In the scanner room, they were given earplugs and MR compatible headphones. Several sensors were fitted on the DA, including a pulsemeter to record cardiac signals; a pneumatic sensor around the waist to measure respiration; and two electrodes on 2 of the left-hand fingers, to record skin conductance. Before placing the coil, DAs were shown the button response box on which they were to provide their answers. They were then moved into the scanner to begin image acquisition. Finally, a laser-safe cloth was draped over the opening of the bore, to ensure that, in case a laser emission went astray, it would not harm the participant.

While DAs were being prepared for scanning, they were led to believe that PRs were undergoing a pain calibration procedure in the MRI control room. In reality, no such procedure was performed, as laser pain stimulation could not be administered in the control room due to safety constraints related to glass walls at the center. Consequently, a deception was introduced whereby DAs believed that PRs would be the recipients of pain on half of the task trials and therefore required calibration. After DAs were positioned in the scanner and the initial MR localizer sequence was launched, PRs were paid 30 CHF, fully debriefed, and dismissed. DAs were informed at the end of the experiment that PRs had not received pain; throughout the task, all DAs believed that a real person was present and potentially affected by their decisions.

##### *MRI acquisition*

Scans were acquired in a 3T Siemens MRI scanner located at the Campus Biotech MRI platform. Sequences during the study were acquired as follows: a field-map scan (2D GR TR=677ms/TE=4.92ms/FA=60deg/SO=PFP); two functional scans of about 20 mins each ([112 112 72], 2D EP TR=1100ms/TE=32ms/FA=50deg, Mosaic Multi-band) where the task was administered; a T1-weighted anatomical scan ( [256 256 208], 3D GR\IR TR=2200ms/TE=2.96ms/FA=9deg/SO=IR\PFP).

##### *Pain Calibration Procedure*

The experimental task required 3 levels of pain (low, medium and high) to be made available to the decision target. Because pain is a subjective experience, we performed a calibration procedure on the DA, following their field map scan acquisition and prior to starting the task. This calibration procedure consisted of stimulating the area around the DA's ankle bone, as this body part was 1) accessible to the experimenter; 2) glabrous skin. Pain stimulations were evoked via nociceptive radiant heat stimulation using an infrared neddymium: yttrium-aluminium-perovskite laser system (Nd: YAP; Stimuli 1340; El. En, Florence; lambda = 1340 nm; pulse duration, 4 ms; beam diameter 6 $mm^2$). The wand of the laser stimulation was passed through an opening in the wall separating control and MR rooms to allow the experimenter to administer stimulations on the DA during functional scans. Stimulations at energies of 1.5-3 J were emitted randomly in step intervals of .25 J, three times each. After each stimulation, DAs rated the pain experienced on a scale presented on the in-bore screen using the button response box. The scale ranged from  to 10, with  indicating no pain, and 1 a first pain, and 10 representing maximum pain, consistent with our previous research(1). Responses for each energy level were then averaged such as to obtain a low, medium and high pain level (average ratings closest to 1; 5 and 9, respectively). In cases where a pain level was equal to another, an additional .25 J was added to the higher pain level. Once individual pain levels were computed, the 3 pain levels were presented to the DA once more, to ensure the participant knew which pain corresponded to their assignation of low, medium and high levels. During the calibration and the task, all parties present were required to wear laser safety goggles. The only exception was the DA whose upper half of the body was protected by laser-safe curtains certified for the wavelength of the stimulator.

##### *Task procedure*

DAs performed two blocks of 18 trials of a decision-making task in the scanner. At each block, participants started off with an endowment of 216 monetary units (MUs), equivalent to 21.6 CHF. At each trial, DAs were first presented with a visual cue, depicting a probability (in the form of a red pie chart on a grey background), and an intensity (represented by 2 or 3 red lightning bolts) of threatened pain, for 6 seconds (Figure 1B). Probabilities ranged from .1-0.9 in steps of .1. DAs were then given 6 seconds to either gamble to avoid the pain altogether, with a 50% chance of winning; or select a sure option, where, if pain were the outcome, it would certainly be reduced by one level. Participants that did not respond in time incurred a penalty of 10 MUs. In the next screen, participants were asked to bid on their decision, selecting a price that ranged from -10 MUs on a scale. Participants that did not respond in the allotted 6 s entered a default "0" for their bid. DA bids were then compared to a hidden reserve price randomly selected for each trial. Bids matching or exceeding this reserve price were accepted as purchase prices for the initial decisions, and the latter were applied to initial values of expected pain, while rejected bids left the DA facing the initial probability and intensity of threatened pain. The bid status (accepted or rejected) was presented on the screen for 6 seconds. Finally, pain was delivered depending on resultant probabilities and intensities. A visual cue representing this pain delivery was simultaneously presented on the in-bore screen, as either initial values of pain intensity (2 or 3 red lightning bolts); reduced values of pain intensity (2 or 3 yellow lightning bolts) or no pain (3 greyed out lightning bolts). In no pain trials, DAs received a non-painful laser stimulation of .5 J. The outcome screen was presented for 6 seconds. Finally, DAs were asked to rate the pain outcome on a visual rating scale, this time anchored by two emojis, one happy and the other unhappy. The scale ranged from 0 to 10, with 10 replaced by the happy emoji. Trials were jittered by an inter-trial interval selected from a 2-5 s range, during which a black fixation cross on a grey screen was presented (Figure 1B). DAs played one block where the decision target was the PR; and another where it was themselves. Block order was randomized across participants, as were buttons assigned to gambles or choices (left and right, to avoid lateralization effects) and placement of happy/unhappy emojis (left/right). Pain was genuinely administered to the DA during DA target blocks, but no actual pain was delivered during PR-target blocks.

Prior to performing the tasks, DAs played 3 practice rounds of the task, with the experimenter guiding them through each screen over the microphone. Only once participants confirmed their understanding of the task did the latter commence. Further, each block began with task instructions presented onscreen. Participants could begin the task when ready, by pressing a button. At the end of the two experimental runs, one payout of the blocks was randomly selected as payment (rounded down or up to the nearest integer) in addition to time paid for participating (50 CHF). Finally, we acquired the T1-MPRAGE anatomical scan, before removing the DA from the scanner. Once out of the scanner, the DA filled out a short series of questionnaires, before being paid, debriefed and sent home.

#### *Computational Models*

To capture the underlying inference hypothesized in this paper, we apply computational models of decision-making under uncertainty to characterize neural processes related to presented stimuli and outcome; and to explain observed behavioral choices. We employ pre-existing models testing the effect of uncertainty on economic decision-making (2), adapted to the domain of pain assessment.

We employed computational models to quantify key variables related to decision-making under pain uncertainty. These variables were calculated on a trial-by-trial basis.

At the initial cue presentation, the expected pain ($EP_1$) represents the average pain intensity given the probabilities of the outcomes. Assuming one outcome is a potential pain intensity $P$, with probability $p_1$, and the other outcome is no pain (intensity):

$$EP_1 = p_1 \cdot P + (1 - p_1) \cdot 0$$

Since the alternative outcome is no pain (intensity 0), the second term vanishes, and this simplifies to:

$$EP_1 = p_1 \cdot P$$

Where:

- $P$: The threatened pain stimulation intensity (e.g., 2 or 3 lightning bolts).
- $p_1$: The initial probability of experiencing pain P (from the visual pie chart).

Initial Risk ($\sigma_1^2$) quantifies the uncertainty (variance) associated with the expected pain at cue presentation. Given the same two outcomes ($P$ with probability $p_1$, and 0 with probability $1-p_1$), the variance is calculated as:

$$\sigma_1^2 = p_1 \cdot P^2 - (EP_1)^2$$

Following the bid outcome, the expected pain ($EP_2$) is updated based on the participant's choice (gamble or sure option) and whether the bid was accepted.

If the gamble option was chosen and the bid was accepted: The probability of pain is halved (to $p_1/2$). Consequently EP and Risk were updated as follows:

$$EP_2 = p_1/2 \cdot P$$

$$\sigma_2^2 = (p_1/2) \cdot P^2 - (EP_2)^2$$

If the sure option was chosen and the bid was accepted: The pain intensity is reduced by one level ($P - 1$), while the probability remains $p_1$. Consequently, EP and Risk were updated as follows:

$$EP_2 = p_1 \cdot (P - 1)$$

$$\sigma_2^2 = p_1 \cdot (P-1)^2-(EP_2)^2$$

If the bid was rejected (for either gamble or sure choice): EP and Risk remained unchanged.

$$EP_2 = EP_1$$

$$\sigma_2^2 = \sigma_1^2$$

Within this framework, Pain Outcome ($P_o$) refers to the actual pain experienced or witnessed at the trial outcome. Instead, Signed Pain Prediction Error (PE) refers the difference between the observed pain outcome and the updated expected pain:

$$PE= P_o - EP_2$$

Finally, we estimated risk prediction error, or "Surprise", as the deviation of the squared signed prediction error from the expected variance after the bid outcome. This is an unsigned error, reflecting the unexpectedness of the outcome's magnitude relative to the updated risk (2).

$$Surprise = PE^2 - \sigma_2^2$$

*Subjective Probability Weighting (Bid Model)*

To assess whether participants distorted objective pain probabilities during decision-making, we fitted a one-parameter Prelec probability weighting function (3) to each participant's bids, separately for self and other conditions. The Prelec function transforms objective probabilities $p$ into subjective decision weights:

$$w(p) = exp(-(-ln\ p)^\wedge\alpha)$$

where $\alpha = 1$ indicates no distortion (objective model), $\alpha < 1$ indicates overweighting of small probabilities (inverse-S shaped distortion), and $\alpha > 1$ indicates overweighting of large probabilities. Using each participant's fitted α, we computed subjective expected pain (sEP = w(p) × Pain) and subjective risk (sRisk = w(p) × $Pain^2$ × (1 − w(p))), as well as subjective prediction error and surprise derived from these quantities.

The bid model assumed that participants' bids were a linear function of subjective expected pain and subjective risk:

$$Bid = \beta_0 + \beta_1 \times sEP + \beta_2 \times sRisk + \varepsilon$$

where $\varepsilon \sim N(0, \sigma^2)$. Five parameters ($\alpha$, $\beta_0$, $\beta_1$, $\beta_2$, $\sigma$) were estimated per participant per target condition by minimizing the negative log-likelihood using unconstrained Nelder-Mead optimization (MATLAB fminsearch) with 20 random restarts. Parameter bounds were enforced via sigmoid ($\alpha \in [0.1, 3.0]$) and exponential ($\sigma > 0$) transforms. A null model fixing $\alpha = 1$ (no probability distortion) was fitted for comparison. Model comparison was performed using AIC. The Prelec function was fitted to bids because they provide a continuous, trial-wise valuation and therefore anchor α in subjective behavior. Two participants (subjects 12 and 22) who placed zero bids on all self-condition trials were excluded from self-condition model fitting (α set to 1.0 for fMRI regressor computation). All other 64 subject-condition fits converged successfully.

#### Data Analysis

##### *Behavioral*

Behavioral data analysis was performed using Matlab 2014b. We analyzed overt, behavioral responses as a function of decision-making variables listed above; and decision targets. We first modelled participants behavioral responses at the level of 1) choice; 2) bids and 3) ratings. We constructed a linear mixed model under the binomial distribution, casting choice (0 = gamble, 1 = sure option) as a function of $EP_1$, $Risk_1$, and choice reaction time (RT), as well as their interactions with the target (Self or Other). In the second, we constructed a general linear model casting bids as a function of $EP_1$, $Risk_1$, choice, and bid RT, as well as their interactions with the target. In the third, we constructed a general linear model casting pain rating as a function of pain, pain prediction error, pain surprise, and rating RT, as well as their interactions with the target. In all models, gender was specified as a nuisance fixed factor, and participants' identifier as a random factor. These models can capture how covert computations relating to uncertainty (expected pain values, pain risk, pain prediction error and pain surprise) may affect overt behavior, and distinguish potential differences in their effects on the target (Supplementary Table S2).

##### *Neuroimaging analysis*

Neuroimaging preprocessing and analysis was performed using SPM12. The main goal of the neuroimaging analysis was to apply a model-based approach to BOLD responses at specific timepoints of the trial that do not require an overt response from the participant. We first describe the preprocessing pipeline applied to the functional MR datasets below.

##### *Preprocessing*

Functional MR volumes were first converted to nifti format from dicom. Images were then slice-time corrected to the middle slice (36). Voxel displacement maps were then applied to the images, before the latter were realigned and unwarped according to head motion parameters (6; 3 translational and 3 rotational). Individual T1w anatomical volumes were then co-registered to the mean functional images and segmented according to six-class tissue probability maps (TPMs). The resulting parameters (forward deformations) were then applied to functional volumes, which were then normalized to the standard MNI152 template, before being smoothed to a FWHM of 8 mm. The resulting preprocessed images were used for analysis.

##### *First Level Model*

In a first instance, we constructed a general linear model including the following onsets: cue presentation (0s); choice response (0s); bid response (0s); bid outcome (0s); trial outcome (0s for self, and 6s for other, respectively); and rating response (0s). These onsets were convolved with the canonical hemodynamic response function and parametrically modulated by the expected value and risk following choice; choice RT; the expected value and risk following bids; bid RT; pain level, pain prediction error (signed PE) and pain surprise (unsigned PE) following trial outcome; and rating and rating RT following pain evaluation. All parametric modulators were serially orthogonalized relative to one another. In addition, we included 24 noise regressors comprised of the 6 motion parameters and 18 cardiac and respiratory regressors (TAPAS toolbox). We employed the FAST algorithm to control for auto-correlation. The general linear model was performed on individual subject timeseries first, before pooling resulting contrasts of interest in a second-level, random-effects model to test group-level responses.

Contrasts of interest focused on the following elements of the computational model, split into predictive and trial outcome phases. In the predictive phase, we tested for expected value of pain and pain risk (concatenating regressors from onsets at cue presentation and following bid outcome). In the trial outcome phase, we tested for pain intensity, signed prediction error and pain surprise (unsigned prediction error).

#### *Group Level Model*

Group-level random-effects analyses were performed using one-sample t-tests on first-level contrast images. Activations were considered significant if exceeding a threshold corresponding to $p < 0.05$ FWE corrected for multiple comparisons at the cluster level, with an underlying voxel-based threshold corresponding to $p < 0.001$ (uncorrected).
To complement the mass voxelwise approach we selected 3 brain regions of theoretical interest: anterior insula, caudate and putamen (dorsal striatum). These regions were derived from the AAL3 atlas (4). The only exception was in the case of prediction error contrasts (signed and Surprise), where the Anterior Insula ROIs were defined functionally from the thresholded masks of surprise-responsive voxels reported in Loued-Khenissi et al., 2020, as an independently defined small volume. Region-of-interest analyses were performed using small volume correction within the bilateral regional masks for expected pain, risk, and pain intensity, and within the surprise region for signed prediction error and surprise. Within these ROIs, an initial voxel-level threshold of $p < 0.005$ (uncorrected) was applied, and effects were considered significant if they survived peak-level FWE correction at $p < 0.05$ for the small volume.

#### *Representational Similarity Analysis*

Representational similarity analysis (RSA) was used to test whether the relational structure of neural activity patterns reflected the relational structure of computational variables across trials. Unlike univariate analyses, which assess whether overall signal amplitude scales with a variable of interest, RSA evaluates whether trials with similar computational properties evoke similar multivoxel activation patterns within a region.

Specifically for each participant, target condition (self, other), and ROI, neural dissimilarity matrices were computed as 1 minus the Pearson product-moment correlation between trial-wise voxel patterns. These dissimilarity matrixes were compared with model-based matrices derived by trial-wise differences in EP, Risk, signed PE, and Surprise. Neural and model-based were compared using Spearman rank correlation $\rho$.

In particular four parameters were collected:

- self-pain similarity: focused only on trials within the self condition.
- other-pain similarity: focused only on trials within the other condition.
- within-target similarity: combining self- and others- trials, and estimating dissimilarity matrices only when comparing trials of the same target
- cross-target similarity: combining self- and others- trials, and estimating dissimilarity matrices only when comparing trials of the different targets

For self-pain and other-pain parameters, group-level inference was assessed using permutation testing (10,000 permutations) to reliably identify $\rho > 0$ correlation, with Benjamini-Hochberg false discovery rate (FDR) correction applied across the regions of interest within each variable. Significant positive correlations indicate that the representational geometry within a region preserves the similarity structure of the corresponding computational variable.

Within- and cross-target similarity parameters allowed us to ascertain whether, and to which extent, the representational code for each parameter was specific/common across targets. Significant evidence of *within-target* $\rho$ > *cross-target* $\rho$, would indicate that regional information encoded is in part target-specific. Instead *cross-target* $\rho > 0$ would reflect the presence of shared code.

#### S2. Prelec probability weighting model

Prelec probability weighting functions were successfully fitted to 64 of 66 subject-condition combinations (31 self, 33 other). Median α values were .818 (self) and .971 (other), indicating modest probability distortion centered near the identity function. The subjective model did not reliably outperform the objective model: the subjective model was favored by AIC in only 9 of 31 self and 12 of 33 other participant fits. Weighting parameters did not differ between pain targets (Wilcoxon signed-rank p = .40). Objective and subjective regressors were highly correlated (EP: median r = .98; Risk: median r = .86), supporting the use of objective task-derived regressors as parametric modulators in the fMRI analysis. The following results are reported using the objective regressors; a supplementary analysis confirmed convergence with subjective regressors for all key findings.

Table S2 reports the fitted Prelec α parameters for each condition.

**Table S2.1** *Prelec α parameter estimates from the bid model.*

| | n fitted | Median α | Mean α | Min | Max | AIC wins |
|---|---|---|---|---|---|---|
| Self | 31 | 0.818 | 1.135 | 0.100 | 3 | 9/31 |
| Other | 33 | 0.971 | 1.456 | 0.100 | 3 | 12/33 |

The Wilcoxon signed-rank test comparing α values between self and other conditions was not significant ($p$ = .42, $n$ = 31 paired), indicating that probability distortion did not differ by target condition.

**Table S2.2** *Per-subject correlations between objective and subjective regressors.*

| Regressor | Target | Median r | Min r | Max r | MAD |
|---|---|---|---|---|---|
| EP | Self | 0.9775 | 0.7018 | 1.0 | 0.2816 |
| EP | Other | 0.9624 | 0.7018 | 1.0 | 0.3390 |
| Risk | Self | 0.8640 | 0.7458 | 1.0 | 0.3071 |
| Risk | Other | 0.8061 | 0.7458 | 0.9997 | 0.4052 |

Expected pain regressors were highly correlated between objective and subjective formulations (median $r > .96$), while risk regressors diverged more substantially (median $r = .81–0.86$), reflecting the nonlinear dependence of variance on the probability weighting function.

A choice-level model was additionally fitted, estimating three parameters per participant per condition: α (Prelec curvature), β (inverse temperature), and θ (risk attitude, where positive values indicate risk aversion). Of 66 participant-condition pairs, 58 had sufficient choice variability for fitting. However, the inverse temperature parameter β converged to near zero for approximately half of participants, rendering the risk attitude parameter θ unidentifiable (values ranged from $-53 \times 10^6$ to $+3.5 \times 10^6$). This reflects the fundamental limitation of fitting three free parameters to 18 binary observations. Among the subset of participants with interpretable fits ($\beta > .5$; approximately 15 participants), θ values clustered in the range .2–1.7, qualitatively consistent with the risk aversion observed in the behavioral GLMEs, but the reduced sample precluded reliable self vs. other comparison.

### S3. Analysis of behavioral measures

Table S3 reports specifications of the (Generalized) Linear Mixed models implemented for the analysis of behavioral responses. As in Model 1 responses were coded by a dichotomous choice variable, a binomial distribution was chosen instead. Models 2-3 were instead analyzed following a normal distribution.

**Table S3** *Each model specifies the outcome variable, primary predictor variables (including the interaction thereof), nuisance variables, and random factors (with random intercept). EP = Expected Pain, RT = Response Times (ms), PE = signed Prediction Error.*

| Model | Dependent Variable | Predictor Variables | Nuisance Variables | Random Factors |
|---|---|---|---|---|
| Model 1 *(binomial)* | Choice *(0 = gamble, 1 = sure option)* | $EP_1$, $Risk_1$, Target, | Sex | Participant |
| Model 2 | Bid *(Monetary Units)* | $EP_1$, $Risk_1$, Bid RT, choice, Target | Sex | Participant |
| Model 3 | Rating [1-10] | Pain, Pain PE, Pain Surprise, choice, Bid, Rating RT, Target | Sex | Participant |

### S4. Full univariate results

This section reports the complete univariate results underlying the main text: small-volume-corrected peaks within a priori regions of interest (Table S4.1) and full voxelwise analysis.

**Table S4.1.** Small-volume-corrected (peak-FWE) responses for pain risk and pain surprise. Surprise was assessed within the *a priori* domain-general surprise region (Loued-Khenissi et al., 2020); Risk and Pain Outcome were assessed within anatomical regions of interest.

| Contrast | Region | peak T | MNI (x, y, z) | p(FWE) |
|---|---|---|---|---|
| Risk, self | L caudate | 3.97 | -16, 18, 12 | .017 |
| Risk, other | L anterior insula | 3.96 | -44, 4, -6 | .025 |
| | R anterior insula | 3.86 | 36, 6, 4 | .031 |
| | L caudate | 4.58 | -14, 2, 18 | .004 |
| | R caudate | 4.69 | 18, 2, 16 | .003 |
| | L putamen | 5.28 | -22, 2, 8 | < .001 |
| | R putamen | 5.37 | 28, -6, -4 | < .001 |
| Risk, other > self | L anterior insula | 4.05 | -44, 4, -8 | .020 |
| | R putamen | 3.71 | 28, -10, -6 | .038 |
| Pain Outcome, other | R anterior insula | 3.86 | 38, 14, 0 | .024 |
| Surprise, self | R anterior insula* | 3.90 | 48, 26, -10 | .016 |
| Surprise, other | R anterior insula* | 3.43 | 44, 26, -2 | .039 |
| *functionally-defined from Loued-Khenissi et al. 2020 | | | | |

**Supplementary Table S4.3** Whole-brain clusters surviving cluster-level FWE correction ($p < 0.05$; cluster-forming threshold $p < 0.001$, uncorrected), objective regressors.

| **Region** | **k** | **p(FWE)** | **T** | **Z** | **x** | **y** | **z** |
|---|---|---|---|---|---|---|---|
| **Parametric effects by target** | | | | | | | |
| **Expected pain — Other** | | | | | | | |
| R occipital pole | 1263 | <0.001 | 7.37 | 5.59 | 18 | -96 | 12 |
| *R superior occipital gyrus* | | | 7.07 | 5.45 | 30 | -92 | 14 |
| R fusiform gyrus | 602 | <0.001 | 7.02 | 5.42 | 28 | -48 | -16 |
| *R occipital fusiform gyrus* | | | 6.67 | 5.24 | 30 | -68 | -10 |
| *R inferior occipital gyrus* | | | 4.20 | 3.72 | 50 | -66 | -6 |
| L occipital fusiform gyrus | 892 | <0.001 | 6.37 | 5.08 | -26 | -80 | -10 |
| *L cerebellum* | | | 5.44 | 4.54 | -30 | -54 | -20 |
| *L occipital fusiform gyrus* | | | 5.06 | 4.30 | -26 | -66 | -12 |
| L middle occipital gyrus | 666 | <0.001 | 5.16 | 4.37 | -24 | -88 | 12 |
| L supplementary motor cortex | 188 | 0.034 | 4.76 | 4.11 | -4 | 14 | 42 |
| *L supplementary motor cortex* | | | 4.33 | 3.81 | -6 | 10 | 56 |
| *L supplementary motor cortex* | | | 3.91 | 3.51 | 0 | 8 | 48 |
| ***Expected pain — Self — no surviving clusters*** | | | | | | | |
| **Risk — Other** | | | | | | | |
| L middle cingulate gyrus | 489 | <0.001 | 5.52 | 4.60 | -6 | 18 | 34 |
| *R middle cingulate gyrus* | | | 4.66 | 4.04 | 10 | 10 | 38 |
| *L supplementary motor cortex* | | | 4.58 | 3.99 | -8 | 12 | 52 |
| R putamen | 533 | <0.001 | 5.37 | 4.50 | 28 | -6 | -4 |
| *R caudate* | | | 5.18 | 4.38 | 16 | 4 | 12 |
| *R amygdala* | | | 5.07 | 4.31 | 24 | -6 | -12 |
| L putamen | 546 | <0.001 | 5.28 | 4.45 | -22 | 2 | 8 |
| *L thalamus* | | | 4.89 | 4.19 | -16 | -20 | 6 |
| *L caudate* | | | 4.58 | 3.98 | -14 | 2 | 18 |
| R superior occipital gyrus | 968 | <0.001 | 5.27 | 4.44 | 26 | -86 | 12 |
| *R calcarine cortex* | | | 4.66 | 4.04 | 30 | -68 | 10 |
| *R calcarine cortex* | | | 4.62 | 4.01 | 18 | -70 | 10 |
| L precentral gyrus | 266 | 0.015 | 5.09 | 4.33 | -54 | 6 | 26 |
| *L precentral gyrus* | | | 4.95 | 4.23 | -40 | 2 | 22 |
| R superior frontal gyrus | 537 | <0.001 | 4.87 | 4.18 | 16 | 4 | 54 |
| *R precentral gyrus* | | | 4.83 | 4.16 | 54 | 6 | 24 |
| *R precentral gyrus* | | | 4.50 | 3.93 | 36 | 0 | 52 |
| L occipital pole | 258 | 0.017 | 4.81 | 4.14 | -12 | -98 | 12 |
| *L middle occipital gyrus* | | | 3.73 | 3.37 | -28 | -92 | 12 |
| *L calcarine cortex* | | | 3.42 | 3.13 | -8 | -90 | 10 |
| R lingual gyrus | 579 | <0.001 | 4.80 | 4.13 | 16 | -82 | -14 |
| *R cerebellum* | | | 4.69 | 4.06 | 28 | -70 | -22 |
| *R cerebellum* | | | 4.13 | 3.67 | 34 | -76 | -24 |
| L cerebellum | 581 | <0.001 | 4.79 | 4.13 | -28 | -66 | -22 |
| *L cerebellum* | | | 4.69 | 4.06 | -20 | -74 | -18 |
| *L cerebellum* | | | 4.31 | 3.80 | -40 | -62 | -22 |

| Region | k | p(FWE) | T | Z | x | y | z |
|---|---|---|---|---|---|---|---|
| R inferior temporal gyrus | 579 | <0.001 | 4.76 | 4.11 | 58 | -58 | -12 |
| *R middle temporal gyrus* | | | 4.43 | 3.89 | 60 | -56 | 0 |
| *R inferior temporal gyrus* | | | 4.10 | 3.65 | 48 | -58 | -8 |
| L superior parietal lobule | 312 | 0.007 | 4.50 | 3.93 | -14 | -68 | 52 |
| *L superior parietal lobule* | | | 3.48 | 3.18 | -16 | -70 | 64 |
| ***Risk — Self — no surviving clusters*** | | | | | | | |
| **Pain outcome — Self** | | | | | | | |
| L middle cingulate gyrus | 7235 | <0.001 | 9.69 | 6.58 | 0 | 8 | 34 |
| *L middle cingulate gyrus* | | | 7.70 | 5.75 | -2 | -10 | 44 |
| *R superior frontal gyrus* | | | 7.43 | 5.62 | 14 | -6 | 68 |
| L anterior insula | 1985 4 | <0.001 | 8.88 | 6.26 | -42 | 6 | -6 |
| *L central operculum* | | | 8.52 | 6.11 | -50 | -4 | 12 |
| *L supramarginal gyrus* | | | 8.15 | 5.96 | -54 | -30 | 30 |
| L cerebellum | 467 | 0.005 | 5.03 | 4.28 | -28 | -34 | -32 |
| *L cerebellum* | | | 4.46 | 3.90 | -24 | -66 | -24 |
| *L cerebellum* | | | 4.45 | 3.90 | -20 | -38 | -48 |
| ***Pain outcome — Other — no surviving clusters*** | | | | | | | |
| ***Pain prediction error — Self — no surviving clusters*** | | | | | | | |
| ***Pain prediction error — Other — no surviving clusters*** | | | | | | | |
| **Surprise — Self** | | | | | | | |
| R postcentral gyrus | 229 | 0.035 | 4.72 | 4.08 | 28 | -38 | 58 |
| *R postcentral gyrus* | | | 4.47 | 3.91 | 30 | -34 | 68 |
| *R postcentral gyrus* | | | 3.99 | 3.57 | 24 | -32 | 62 |
| ***Surprise — Other — no surviving clusters*** | | | | | | | |
| **Ratings — Self** | | | | | | | |
| R precuneus | 285 | 0.026 | 5.11 | 4.34 | 10 | -68 | 54 |
| *R superior parietal lobule* | | | 4.38 | 3.85 | 20 | -64 | 50 |
| *R superior parietal lobule* | | | 3.79 | 3.42 | 20 | -68 | 64 |
| ***Ratings — Other — no surviving clusters*** | | | | | | | |
| **Self vs. Other contrasts** | | | | | | | |
| **Pain outcome — Self > Other** | | | | | | | |
| R middle cingulate gyrus | 6905 | <0.001 | 9.39 | 6.46 | 2 | 4 | 38 |
| *L middle cingulate gyrus* | | | 7.81 | 5.80 | 0 | -8 | 44 |
| *L precentral gyrus* | | | 7.20 | 5.51 | -8 | -16 | 74 |
| L central operculum | 8332 | <0.001 | 8.68 | 6.18 | -50 | -4 | 12 |
| *L supramarginal gyrus* | | | 8.10 | 5.93 | -58 | -28 | 26 |
| *L anterior insula* | | | 7.90 | 5.84 | -42 | 6 | -6 |
| R parietal operculum | 6834 | <0.001 | 7.56 | 5.69 | 58 | -28 | 26 |
| *R anterior insula* | | | 7.26 | 5.54 | 36 | 8 | 8 |
| *R anterior insula* | | | 6.91 | 5.36 | 42 | -4 | 4 |
| brainstem | 2798 | <0.001 | 6.67 | 5.24 | 6 | -36 | -2 |
| *R cerebellum* | | | 6.24 | 5.01 | 20 | -46 | -48 |
| *brainstem* | | | 5.40 | 4.52 | -4 | -38 | -4 |
| **Surprise — Self > Other** | | | | | | | |
| R postcentral gyrus | 260 | 0.023 | 5.03 | 4.29 | 28 | -38 | 58 |

| Region | k | p(FWE) | T | Z | x | y | z |
|---|---|---|---|---|---|---|---|
| *R postcentral gyrus* | | | 3.95 | 3.54 | 24 | -30 | 66 |
| *R postcentral gyrus* | | | 3.42 | 3.14 | 34 | -28 | 54 |
| **Expected pain — Other > Self** | | | | | | | |
| L fusiform gyrus | 229 | 0.026 | 5.12 | 4.35 | -24 | -54 | -14 |
| *L lingual gyrus* | | | 3.52 | 3.21 | -28 | -56 | -6 |
| **Risk — Other > Self** | | | | | | | |
| R middle temporal gyrus | 575 | <0.001 | 5.35 | 4.49 | 64 | -48 | 2 |
| *R middle temporal gyrus* | | | 4.92 | 4.21 | 60 | -56 | 0 |
| *R middle temporal gyrus* | | | 4.91 | 4.21 | 64 | -56 | -8 |

Clusters significant at cluster-level familywise-error (FWE) $p < 0.05$, cluster-forming (voxel-height) threshold $p < 0.001$ uncorrected. For each cluster the peak and up to two further local maxima (> 8 mm apart) are listed; cluster extent (k, voxels) and cluster-level p(FWE) appear on the peak row. Coordinates are MNI (mm); T and Z are the statistics at the maximum. One-sample t-tests across N = 33 (df = 32).
